# Resource abundance and dietary specialization predict elevational migration in a hyperdiverse montane bird community

**DOI:** 10.64898/2026.03.18.710293

**Authors:** Tarun Menon, Abhinav Tyagi, Shreyas Managave, Uma Ramakrishnan, Umesh Srinivasan

## Abstract

Migration is a well-described behavioural strategy that allows species to track variation in resources and climatic conditions by moving in response to seasonality. A common form is elevational migration, an annual short-distance movement undertaken by many mountain bird species globally. While studies show that the timing of migration may relate to food availability, the mechanisms determining which species migrate remain unclear. Our study investigated if the degree of dietary specialization explains why some high-elevation bird species in seasonal environments migrate downslope for the winter while others remain resident at high altitudes despite the apparent scarcity of their preferred food resources. We mist-netted birds along a 2300-m elevational gradient in the Eastern Himalaya and collected blood and faecal samples from 261 individual birds belonging to 18 species of high-elevation residents (ten) and elevational migrants (eight) in their breeding and wintering ranges. Using stable isotope ratios of carbon and nitrogen in whole blood and faecal DNA metabarcoding, we compared their seasonal trophic levels and dietary niches. Nitrogen isotope ratios showed that residents had a substantially lower trophic position in the winter compared to summer (-0.35 [-0.52, -0.17]), whereas migrants had a slightly higher trophic position in the winter (0.15 [-0.02, 0.32]). This trophic shift in residents was likely due to a decrease in insectivory and an increase in frugivory in the winter. The frequency of key insect orders (Lepidoptera, Hemiptera, and Coleoptera) declined by 20-35% in their winter diets alongside an increase in fruit, particularly from the family Polygonaceae (0.33 [0.18, 0.46]). Additionally, compared with residents, migrants showed greater overlap in their dietary niches between summer and winter (98% vs 80%). Because arthropod abundances in the Himalayas peak at high elevations in the summer and decline in the winter, we suggest that elevational migrants are likely dietary specialists that track resources, while high-elevation residents are dietary generalists that supplement their winter diet with fruit and nectar because of the scarcity of arthropods. These findings indicate that a species’ dietary specialization is linked to its migratory behaviour, providing a potential mechanistic explanation for how different species solve the challenge of seasonal resource limitation.

## Introduction

Animal migrations are some of the most spectacular phenomena in the natural world. The seasonal migrations of vast herds of wildebeest for instance, are amongst the most iconic spectacles in nature. While such long-distance movements across continents are well studied, many animals undertake shorter, less visible migrations that are equally remarkable and receive less attention (Hsiung et al. 2018). Elevational migration is a type of short-distance migration defined by the seasonal movement of montane populations between breeding grounds and wintering grounds at different elevations. In most cases, migrants breed at high elevations and winter at lower elevations (with some exceptions of birds moving downslope to breed; Hess et al. 2012, Tsai et al. 2021). Amongst montane species, elevational migration is a common phenomenon both taxonomically and geographically (Boyle 2017, Hsiung et al. 2018); however, birds are the most conspicuous and well-documented taxon that migrate elevationally, accounting for over 60% of studies on the phenomenon (Hsiung et al. 2018). Across continents, a substantial proportion of montane bird species (ranging from 20% to 70%) engage in seasonal elevational movements (Boyle 2017).

The drivers of migration are widely hypothesized to include a combination of climate-related factors and seasonal shifts in resource availability. In the Neotropics for instance, frugivores migrate downslope to avoid extreme rainfall at higher elevations and back upslope to track fruit abundance (Loiselle and Blake 1991, Boyle et al. 2010). While elevational migration in Neotropical frugivores has been relatively well studied, comparable insights from other regions remain limited (Barçante et al. 2017). In subtropical mountain ranges such as the Himalayas, most elevational migrants are insectivorous (Barçante et al. 2017), and their movements are likely driven by cold temperature-related constraints such as reduced arthropod availability and physiological limitations on thermoregulation. Many studies have shown that the timing of elevational migration is related to food abundance (Loiselle and Blake 1991, Solórzano et al. 2000, Chaves-Campos 2004). However, few have directly tested whether dietary flexibility helps birds respond to seasonal changes in food availability and whether this flexibility might drive their movements. This leaves a gap in our understanding of how food dynamics might shape migratory behaviour.

Arthropods are an important dietary resource for a significant fraction of the world’s birds. Arthropods are also particularly sensitive to seasonal changes in temperature because they are small ectotherms (Thomas et al. 1994). This results in arthropod abundances dropping significantly at high elevations, while low elevations have a relatively higher number of arthropods in the winter (Menon 2026). The relatively short growing season at high elevations results in a synchronous emergence and reproduction of arthropods, causing a sharp summer peak in their abundance (Supriya et al. 2019). The lack of arthropods at high elevations in winter might drive insectivorous bird species that breed at high elevations to migrate downslope in the winter to access food and then move back up in the summer to breed during the resource-rich arthropod pulse. However, a significant proportion of high-elevation bird species that feed predominantly on arthropods in summer continue to overwinter at high elevations despite the relative lack of arthropod resources.

In this study we focus on the food availability hypothesis of elevational migration and examine if dietary specialization potentially explains “why some, but not all, bird species migrate” (Fretwell 1980). Dietary specialization might indeed play an important role in driving movement; for instance, among closely related frugivores in Costa Rica, migrants were found to have stronger preferences for fruit than their resident counterparts (Boyle et al. 2011). If migratory species show a similar stronger preference for arthropods (over fruit and nectar), we hypothesize that they would be dietary specialists that track arthropod availability along the elevation gradient across seasons. We further hypothesize that high-elevation resident (i.e., non-migratory) species, on the other hand, are dietary generalists that, when faced with a reduction in preferred arthropod availability in winter, supplement their diet with other available resources such as fruit and nectar. Indeed, despite the harsh winter climate, several high-altitude plant taxa in the Eastern Himalaya, most notably the genera *Polygonum* (knotweed), *Ligustrum* (privet) and *Leycesteria* (honeysuckle) were observed by the authors to fruit abundantly during the winter months, a pattern corroborated by regional phenological studies (Omori and Ohba 1996) and flora (Sankara Rao and Kumar 2026). Alternatively, residents could also be dietary specialists that have more specialized foraging strategies allowing them to access dormant arthropods or those surviving within more buffered microhabitats such as within tree bark and rock crevices. We acknowledge that these observed patterns could potentially arise independent of food limitation; if migration is driven by climate tracking, migrants and residents might simply consume the most readily available resources at a given elevation. However, climatic constraint hypotheses are insufficient to explain elevational migration as studies show that migratory species do not consistently track their thermal niches across seasons (Menon et al. 2023, Somveille et al. 2026).

To test these hypotheses, we characterised and compared the diets of migratory and resident species in their breeding and non-breeding seasons in the eastern Himalayas. We used a combination of stable isotope analysis and faecal DNA metabarcoding to provide a complimentary and comprehensive understanding of species’ dietary niches (Hoenig et al. 2022). We used stable isotope ratios of nitrogen (δ^15^ N) and carbon (δ^13^C) in whole blood which represents diet assimilated over an approximately two-week period (Ogden et al. 2004). Because the heavier isotope of nitrogen (^15^N) gets enriched in the tissues of consumers, higher values of δ^15^ N represent a higher trophic level and, in this case, greater insectivory (Newsome et al. 2007). Analysing variability in δ^15^ N and δ^13^ C in bivariate space allows us to understand an individual bird’s isotopic niche, which is closely related to its dietary niche (Newsome et al. 2007, Jackson et al. 2011). Faecal DNA metabarcoding, on the other hand, allows us to identify all the dietary components with greater taxonomic resolution and understand how diet may change across seasons. Combining the two methods allows us to detect trophic shifts while simultaneously identifying the taxonomic groups that contribute to these shifts. We make two key predictions; 1) elevational migrants are likely to have a similar trophic level in both seasons and will have greater overlap in their isotopic niche and dietary community composition across seasons, and 2) high-elevation residents will have a lower trophic level in the winter because of lower arthropod and higher fruit/nectar in their winter diets and this will also lead to relatively lesser overlap in their isotopic niche and dietary community composition across season. However, if residents can access arthropods even in the winter arising from specialized foraging strategies, they will have similar trophic levels and dietary niches (greater overlap) in either season which matches the patterns observed in migrants.

## Methods

### Study Site

We conducted this study in Eaglenest Wildlife Sanctuary (EWS) and the adjoining Singchung Bugun Village Community Reserve (SBVCR) in Arunachal Pradesh, India. This landscape is part of the Eastern Himalaya Global Biodiversity Hotspot and home to >350 breeding bird species, where over 65% of high elevation breeders migrate elevationally (Myers et al. 2000, Menon et al. 2025). EWS covers an area of 217 km^2^ with an elevation gradient ranging from 500 m to 3250 m above sea level (ASL). The low elevations (below 1000 m ASL) are home to tropical wet evergreen forests transitioning to broadleaved subtropical forests at ∼1800 m ASL. Broadleaved temperate forests consisting of oak and rhododendron can be found between 1800-2800 m and above 2800 m, the main habitat is coniferous temperate forests.

### Field Sampling

Bird species at our field site usually establish their breeding territories by early April and reach their wintering grounds by early November. Field work was therefore carried out over two breeding seasons (April - June) and two non-breeding seasons (November - January) between 2021 and 2024. In each season we set up 20 mist nets (12 m, four shelf, 16 mm mesh size) in the understorey at seven sites along the elevation gradient at 800 m, 1200 m, 1600 m, 2000 m, 2400 m, 2800 m, and 3100 m ASL. Species that were caught above 2000 m ASL in both the breeding and non-breeding season were considered high-elevation residents while species that were caught above 2000 m ASL in the breeding season and below 1600 m ASL in the non-breeding season were considered elevational migrants. Every bird captured was banded with a uniquely numbered aluminium band and blood and faecal samples were collected from high-elevation residents and elevational migrants (Appendix S1). We additionally collected 74 pairs of leaves of the three most common plant species from every 100 m elevational band (from 800 to 3100m ASL) to establish an isotopic baseline. Since leaf δ^15^ N and δ^13^C do not generally vary with season (Ometto et al. 2006), all leaf samples were collected in the summer of 2022.

### Stable Isotope Methods

We processed blood samples from bird species caught in the latter half of each season to ensure the isotopic signature represents the elevation the bird was caught. The blood samples were dried in a hot air oven at 56°C for 48 hrs. The dried blood sample was then powdered, weighed (∼0.2 mg) and packed in a tin capsule. Plant samples were similarly oven dried, powdered in a mixer mill, weighed (∼1 mg) and packed. The prepared samples were analysed for δ^15^ N and δ^13^C at the Stable Isotope Facility in the Indian Institute of Science Education and Research Pune (IISER) and the Indian Institute of Science Bangalore (IISc). Isotope ratios were expressed in parts per thousand deviations from each standard (‰) which were Vienna PeeDee Belemnite (VPDB) and atmospheric air for δ^15^N and δ^13^C, respectively. The samples were measured together with laboratory standards to ensure repeatability and accuracy; the precision (1-sigma) on measurements at the two facilities was 0.07‰ (IISER) and 0.15‰ (IISc) for δ^15^N and 0.05‰ for δ^13^C. Trophic position (TP) was calculated as:

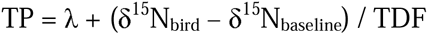

λ represents the estimated TP of baseline producers (λ = 1), δ^15^N_bird_ represents the measured δ^15^N value of the individual bird, δ^15^N_baseline_ represents the mean δ^15^N of plants at low (500-1400 m), medium (1400-2200) and high elevation (>2200 m) and TDF represents the Trophic Discrimination Factor which we considered to be 2.3 ± 0.3 (SD) based on published values for whole blood from an omnivorous bird (Stephens et al. 2023). We estimated TP of all resident and migrant species combined in each season using a Bayesian phylogenetic multilevel model. Standard errors for TP were calculated using first-order Taylor series expansion for error propagation, accounting for variance in δ^15^N_baseline_ and TDF during the calculation of TP. We evaluated the interactive effects of migratory status and seasonality on TP using the brms package (Bürkner et al. 2025) in R (R Core Team. 2021). We treated the phylogenetic lineage of each species as a random intercept to control for evolutionary non-independence, incorporating a phylogenetic correlation matrix derived from the time-calibrated tree by Claramunt et al. (2025). We used weakly informative priors, and the model was run with four Markov Chain Monte Carlo (MCMC) chains for 4,000 iterations each, with a warm-up period of 2,000 iterations. We assessed chain convergence using the R̂ diagnostic, ensuring all values were < 1.01, and by visual inspection of trace plots. In addition to the global model, we ran separate models for each species to estimate species-specific seasonal shifts. Results are presented as posterior means with 95% Credible Intervals [CI].

To visualise isotopic niche, we created biplots of baseline corrected δ^15^N and δ^13^C because stable isotope ratios varied with elevation (Appendix S1: Table S1). Trophic position represents baseline corrected δ^15^ N and baseline corrected δ^13^ C (δ^13^ Ccorr) was calculated using a formula described by Olsson et al. (2009) (Appendix S1). To visualise niche width and overlap we plotted standard ellipses corrected for small sample sizes (SEAc) using SIBER in R (Jackson et al. 2011). For further analysis, we computed Bayesian niche overlap probabilities between the summer and winter isotopic niches of resident and migrant species using the package nicheROVER in R (Swanson et al. 2015). For a given species we plot the niche region where there is a 95% probability of finding an individual in either season. The seasonal niche overlap for that species is then expressed as the probability of finding an individual in the winter within the species’ summer isotopic niche at an alpha of 0.95.

### DNA metabarcoding

DNA from faecal samples were extracted following the manufacturer’s recommendations using the QIAmp Fast DNA Stool Mini Kit (QIAGEN, Germany). Arthropod and plant DNA in each sample was amplified in two separate PCR reactions (Appendix S1) and amplification was checked on 2% agarose gel. We used the ZBJ primer pair (ZBJ-ArtF1c and ZBJ-ArtR2c) that targets the hypervariable region of the arthropod mitochondrial cytochrome c oxidase I (COI) gene (Zeale et al. 2011). To amplify plant DNA we used the trnL primer pair (g and h) that targets the P6-loop of the chloroplast trnL (UAA) intron (Taberlet et al. 2007). PCR products were purified, indexed and sequenced at the NGS facility at the National Centre for Biological Sciences, Bangalore. To ensure the reliability of our sequencing results we also sequenced a number of positive and negative controls. Negative controls were included at the extraction, PCR and indexing stage. We had two positive controls, one consisting of a known mixture of five arthropod orders from our field site and second consisting of five common plant orders. For more details on the primers, PCR conditions, library preparation and sequencing, refer to the supporting information (Appendix S1).

The sequencing reads were demultiplexed and the primer sequences along with the heterogeneity spacers were trimmed. We used the DADA2 pipeline (Callahan et al. 2016) to denoise and dereplicate the remaining reads, remove chimaeric sequences and create an Amplicon Sequence Variant (ASV) table. Contamination was low, read counts lower or equal to the count of an ASV appearing in the negative control was replaced with zero. ASVs with less than 20 sequencing reads across all samples were removed. We assigned taxonomy (arthropods – order level, plants – family level) to the resulting ASVs using ecotag (OBITools; Boyer et al. 2016) and a reference database in ecoPCR format (Appendix S1). ASVs that were less than a 90% match were excluded from further analysis. We also manually removed problematic sequences of plants and arthropods that are unlikely to be found in the region and non-fruiting/flowering plants that may represent secondary consumption (da Silva et al. 2019). The ASV read counts were transformed to their presence or absence in each sample. Since these are primarily insectivorous birds, plants have a higher possibility of being picked up in their diet due to secondary consumption, so low frequency plant ASV reads (<5% of total reads in a sample) were replaced with 0 whereas arthropod ASVs with less than 10 reads in a sample were considered to be absent.

We summarized the seasonal diet composition of residents and migrants (pooled and at the individual species level) using the frequency of occurrence (FOO) of different arthropod orders and plant families in the individual faecal samples. We interpreted higher FOO values as an indicator of greater dietary importance, suggesting that these taxa are more consistently consumed across the sampled individuals. We chose FOO over Relative Read Abundance (RRA) as it offers a conservative measure, mitigating the risk of sequencing or digestion biases misrepresenting the true importance of specific prey. (Lamb et al. 2019). We tested for a seasonal difference in FOO of each major arthropod order and plant family in the diets of residents and migrants (pooled) using a Bayesian phylogenetic logistic regression:

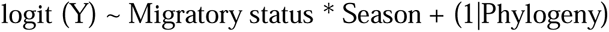

where Y represents the FOO an individual arthropod order or plant family. We applied the same Bayesian framework and sampling parameters used in our trophic position analysis to the models of arthropod and plant occurrence. To analyse seasonal differences in diet composition in terms of the arthropod and plant orders, we performed principal coordinate analysis (PCoA), permutational multivariate analysis (PERMANOVA) and permutational multivariate analysis of dispersion (PERMDISP) using the Jaccard index. We conducted these analyses both for pooled groups (residents and migrants) and individual species. In the pooled models, we accounted for interspecific variation by restricting permutations within species blocks, ensuring that seasonal comparisons were controlled for species-level patterns. All analysis was done using the “vegan” package in R (Oksanen et al. 2007).

## Results

In 10-12 mist net days per site across two years (74 days), we captured 1971 unique individuals belonging to 160 species in summer. In 12 mist net days per site across two years (84 days), we captured 1721 unique individuals belonging to 133 species in winter. We collected blood and faecal samples from all high-elevation resident and migrant species that were reasonably common and were expected to be captured at least ten times in either season over two years. We recovered stable isotope ratios from the whole blood of 112 individuals belonging to seven resident species and 124 individuals belonging to six migratory species. We were able to extract, amplify and successfully sequence 121 faecal samples belonging to ten resident species and 140 samples from eight migrant species. The species were all primarily insectivorous and inhabiting the understory of dense broadleaf forests (Appendix S1: Table S2). High-elevation residents belonged to 8 different families which included both small and large bodied species ranging from the Black-browed Tit *Aegithalos iouschistos* (∼7 g) to the Black-faced Laughingthrush *Trochalopteron affine* (∼70 g). The migrants represented four distinct families, including four species within the family Muscicapidae (Old World flycatchers); all were small-to-medium-sized species weighing less than 30 g.

### Stable Isotope Analysis: Trophic Level

Overall, δ^13^C and δ^15^N values in avian whole blood ranged between -27.5 to -22.8 ‰ and 0.4 to 9.2 ‰ respectively. To test the prediction that residents have a lower-trophic position in winter while migrants maintain their trophic position by tracking arthropods, we used a Bayesian phylogenetic model to analyse isotopic data. The model revealed a distinct interaction between migratory status and season (β_interaction_ = -0.49 [-0.73, -0.26]) indicating that the seasonal shift in trophic position differed between residents and migrants. We found that high-elevation residents had a lower trophic position in the winter compared to the summer with a mean decline of -0.35 [-0.52, -0.17] (Figure 1). Elevational migrants had a relatively stable albeit slightly higher mean trophic position in the winter compared with the summer (0.15 [-0.02, 0.32]) but the 95% CI overlapped zero (Figure 1). We analysed seasonal trophic positions of five resident and six migrant species individually which had at least five samples in either season. The individual species models highlighted variation in the degree of trophic position shifts of individual resident and migrant species across season. We find that all five high-elevation residents have a lower trophic position in the winter, although the magnitude of this shift varied by species (Figure 1; Appendix S1: Table S3). Stripe-throated Yuhina *Yuhina gularis* showed the most pronounced winter decline, with the posterior mean TP dropping from 3.69 [3.40, 3.93] in summer to 2.94 [2.69, 3.20] in winter. Brown-throated Fulvetta *Fulvetta ludlowi* remained relatively stable across both seasons with overlapping credible intervals (mean difference = -0.09 [-0.45, 0.29]; Figure 1). Most migrant species had a relatively stable seasonal trophic position with two species exhibiting a notable increase in estimated trophic position in the winter although 95% CI overlapped zero (Figure 1; Appendix S1: Table S3). The trophic position of Whistler’s Warbler *Phylloscopus whistleri* increased by 0.42 [-0.02, 0.84] and Chestnut-headed Tesia *Cettia castaneocoronata* increased by 0.39 [-0.10, 0.84].

**Figure 1:**
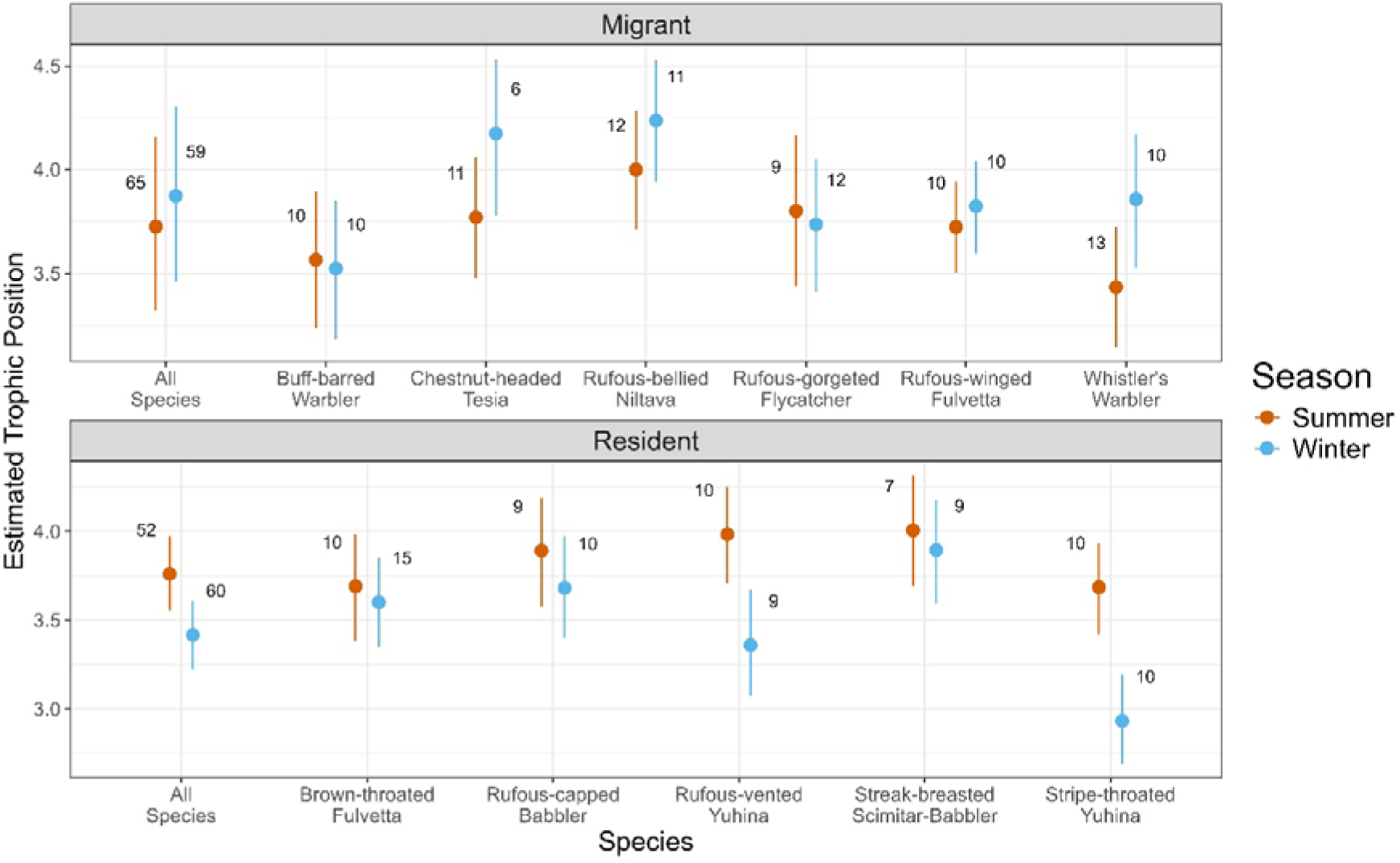
Dot plots of mean posterior trophic position of high-elevation residents and elevational migrants in the summer and winter using a Bayesian phylogenetic multilevel model. The error bars represent 95% credible intervals. Sample sizes in the summer/winter are given next to each plot

### Stable Isotope Analysis: Isotopic Niche

We evaluated the prediction that migrants would exhibit greater seasonal isotopic niche overlap compared to residents by calculating Bayesian niche overlap probabilities. Supporting this prediction, all elevational migrants pooled showed a high median probability (98%) that their summer isotopic niche overlapped with their winter niche (Figure 2). Conversely, pooled residents showed significantly lower seasonal overlap (80%). Among individual species most residents (apart from Streak-breasted Scimitar-Babbler *Pomatorhinus ruficollis*) had little overlap between their summer and winter isotopic niches (Figure 2). Median overlap probabilities were below 50% for most species but was 88% for Streak-breasted Scimitar-Babbler (Appendix S1: Table S4). Most migrants had substantial overlap between their summer and winter isotopic niches (Figure 2). Median overlap probabilities were above 75% for most species but was 58% for Whistler’s Warbler (Appendix S1: Table S4)

**Figure 2:**
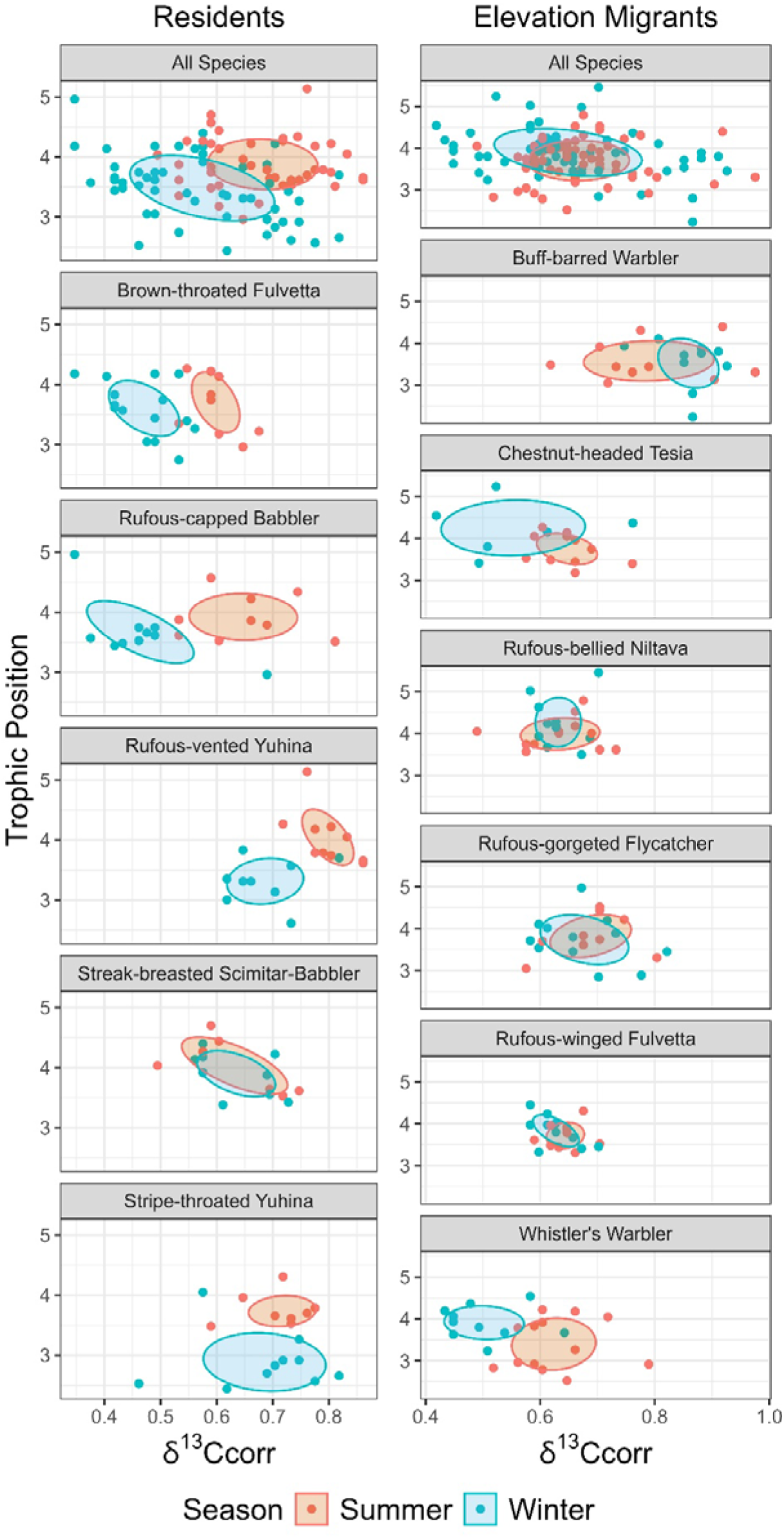
Biplots of baseline corrected δ^15^ N (trophic position) and δ^13^C of high-elevation residents and migrants in the summer and winter. Ellipses represent Standard error ellipses.

### Faecal DNA metabarcoding: Frequency of Occurence

Combining both sequencing runs we generated a total of 60,577,757 paired-end reads (MiSeq: 31,817,757 and NovaSeq: 28,760,000) of 283 samples, which included six negative controls and four positive controls. After merging the paired-end reads and sequence processing, we recovered 5.5 million reads of 3009 arthropod ASVs belonging to 24 orders. Five orders which include Lepidoptera, Diptera, Coleoptera, Hemiptera and Araneae, represent over 90% of all reads. We recovered 14.5 million reads of 420 plant ASVs belonging to 102 families (40 orders) with Polygonaceae and Ericaceae representing over 40% of all reads.

To evaluate the taxonomic basis of these shifts, we used Bayesian phylogenetic logistic regression. We tested our prediction that residents would show a seasonal decline in the frequency of occurrence (FOO) of key arthropod orders alongside an increase in certain plant families, whereas elevational migrants would show stable dietary patterns across seasons (Figure 3; Appendix S1: Table S5). Consistent with this, resident diets (pooled) showed a significant winter reduction in the FOO of Hemiptera (mean difference = -0.18 [95% CI: - 0.33, -0.02]), Diptera (-0.15 [-0.28, -0.01]), Coleoptera (-0.20 [-0.37, -0.02]), and Lepidoptera (-0.35 [-0.47, -0.23]), while Araneae remained stable (-0.05 [-0.22, 0.13]). We analysed seasonal change in the FOO of these arthropod orders in the diets of five resident and six migrant species individually which had at least five samples in either season. All individual resident species show a winter decline in FOO in at least three of these four orders (Appendix S1: Figure S1). Migrant diets (pooled) showed no significant seasonal change in the FOO of Hemiptera (-0.08 [-0.06, 0.21]), Coleoptera (-0.02 [-0.17, 0.13]), or Lepidoptera (-0.02 [-0.05, 0.08]). Notably, while the FOO of Diptera slightly decreased in migrants (-0.1 [-0.2, -0.03]), their consumption of Araneae increased significantly in the winter (0.19 [0.03, 0.35]). Most individual migrant species (except Rufous-gorgeted Flycatcher) showed greater FOO of Araneae in their winter diets while only Rufous-bellied Niltava *Niltava sundara* and Rufous-winged Fulvetta *Schoeniparus castaneceps* showed lower FOO of Diptera and in their winter diets (Appendix S1: Figure S1). With respect to plant families consumed, residents (pooled) showed a significant winter increase in the FOO of Polygonaceae (mean difference = 0.33 [0.18, 0.46]), whereas migrants (pooled) showed a decrease (-0.15 [-0.27, -0.01]). Among individual resident species, the FOO of Polygonacae was highest in the diets of Streak-breasted Scimitar-Babbler (0.75), Rufous-capped Babbler *Cyanoderma ruficeps* (0.64), Brown-throated Fulvetta (0.43) and second highest in Rufous-vented Yuhina *Yuhina occipitalis* (0.37) and Stripe-throated Yuhina (0.16). The families with the highest FOO in the winter diets of Stripe-throated Yuhina were Caprifoliaceae (0.50) and Oleacae (0.50) while in Rufous-vented Yuhina it was Capparacae (0.45) and Rosaceae (0.45). In the summer Ericaceae had the highest frequency of occurrence in all species of residents and migrants except the Rufous-capped Babbler. There were no plant families with more than a 0.2 FOO in the diets of all migrants (pooled) in the winter, nor were there any families consistently seen across all species (Appendix S1: Table S6).

**Figure 3:**
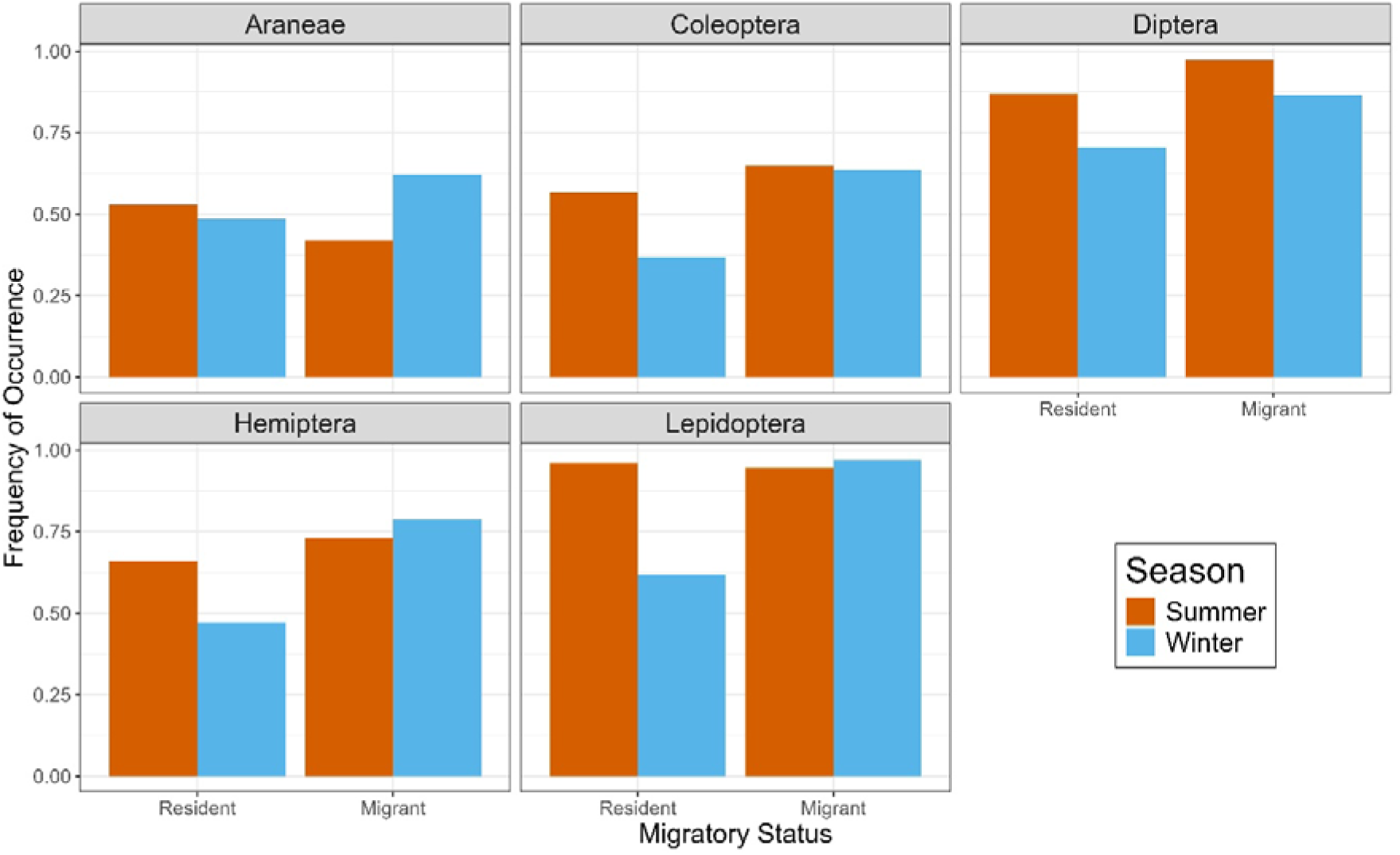
Bar plots comparing the frequency of occurrence of the five most consumed arthropod orders in the diets of high-elevation residents and elevational migrants across seasons.

### Faecal DNA metabarcoding: Diet Composition

The Principal Coordinate Analysis, consistent with our prediction, demonstrates how pooled migrants show greater seasonal overlap in dietary composition compared to pooled residents (Figure 4). We expected PERMANOVA to show significant seasonal differences in dietary composition in residents and migrants however it was significantly different for both with the difference being larger in residents (Resident: Pseudo-F_1,106_ = 7.91, p < 0.001; Migrant: Pseudo-F_1,129_ = 5.76, p < 0.001). Apart from diet composition changing, we expected residents to have a more variable diet in the winter due to the apparent absence of preferred prey which may not be the case with migrants. As expected, PERMDISP showed significant seasonal differences in multivariate dispersion in residents but not migrants (Resident: F_1,106_ = 26.10, p < 0.001; Migrant: F_1,129_ = 2.30, p = 0.13). The dietary compositions of individual migrant species are clustered together while residents like Streak-breasted Scimitar Babbler, Rufous-vented and Stripe-throated Yuhina have very different dietary compositions across seasons (Appendix S1: Figure S2). All resident (except Brown-throated Fulvetta) and migrant (except Rufous-gorgeted Flycatcher) species have significant differences in diet composition across seasons (Appendix S1: Table S7). Three resident (Brown-throated Fulvetta, Rufous-vented and Stripe-throated Yuhina) and zero migrant species showed significant seasonal differences in multivariate dispersion (Appendix S1: Table S8).

**Figure 4:**
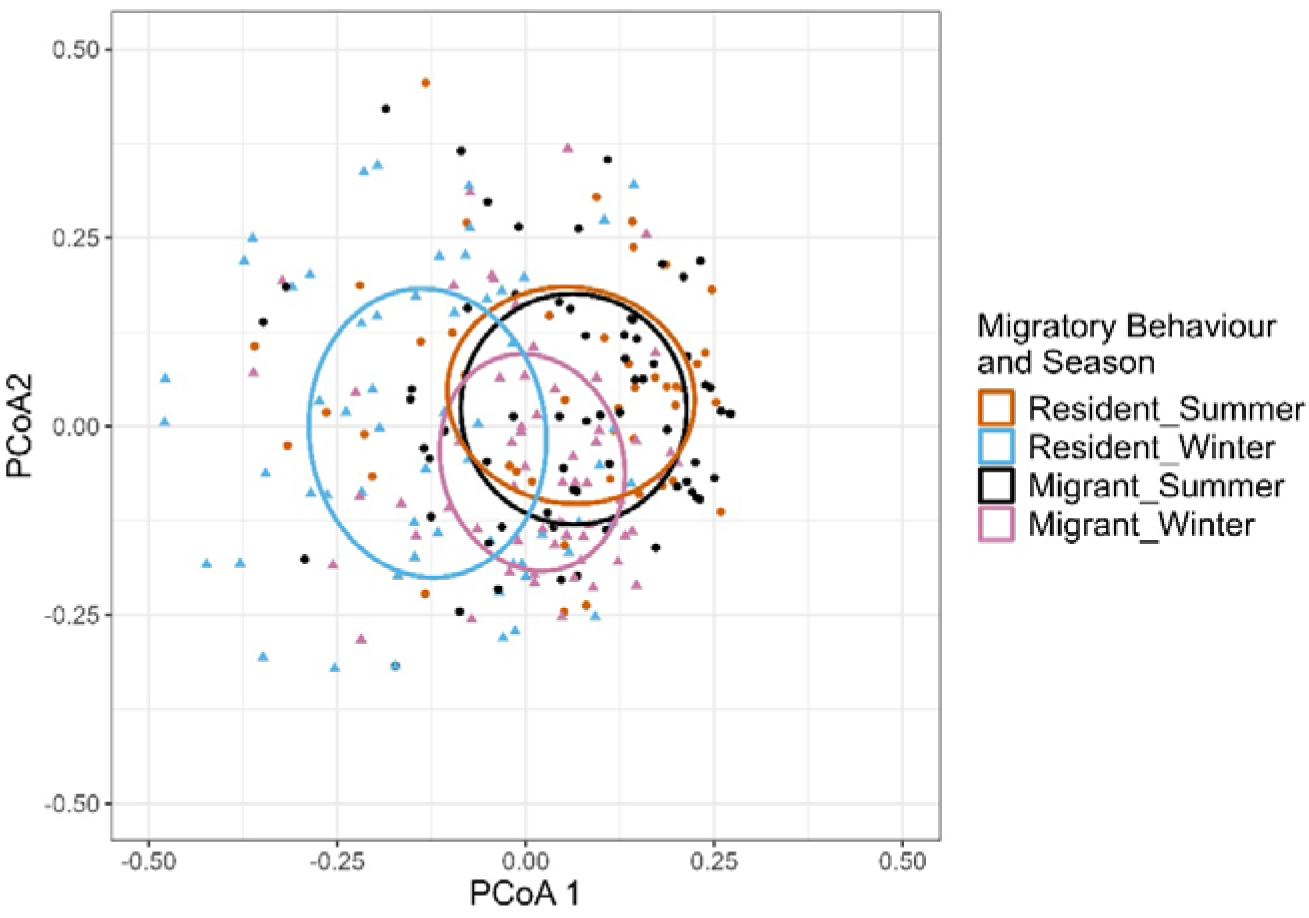
Principal Coordinate Analysis of differences in dietary composition of high-elevation residents and elevational migrants across seasons. Each point represents dietary composition of individual birds in summer (circles) and winter (triangles), and the colors represent season and if the bird was resident or migrant. Ellipses denote the standard error of the centroid diet of residents and migrants in either season at 95% confidence.

## Discussion

Within the larger framework of food limitation as a potential driver of elevational migration, we evaluated how differences in dietary specialization may explain the variation in migratory behaviour in birds. Our results reveal that high-elevation residents are likely dietary generalists that switch from a primarily arthropod-based summer diet to include more plant material in winter, whereas elevational migrants maintain relatively consistent diets across seasons and are likely dietary specialists tracking arthropod resources across elevation and seasons. By combining stable isotope analysis and DNA metabarcoding, we demonstrate how these two methods converge to explain how different species might solve the challenge of seasonal resource limitation.

### Dietary Specialization and resource tracking in migrants

Elevational migrants maintained a relatively stable trophic position across seasons with a slight increase in winter which suggests consistent insectivory. Correspondingly, they have similar frequencies of major arthropod orders such as Hemiptera, Coleoptera, or Lepidoptera in their diets across seasons. Migrants had considerable seasonal overlap in their isotopic niche; however, they had significant differences in dietary compositions with little variation in dispersion across seasons. The difference in dietary composition without a change in isotopic niches may be due to migrants shifting their diets between functionally similar taxa based on the seasonal turnover in arthropod and plant availability at low elevations. The higher trophic positions of migrants in the winter may be driven by an increase in the consumption of spiders which may be attributed to the predatory nature of spiders (higher trophic level) and low elevations having higher spider abundances in the winter (Menon 2026). The two species (Whistler’s Warbler and Chestnut-headed Tesia *Cettia castaneocoronata*) which had a substantially higher trophic position in the winter (Figure 1) also more frequently consumed spiders in the winter compared to the summer (Appendix S1: Figure S1). This highlights the importance of spiders as a winter food source, supporting multiple studies that show spiders to persist through winter due to cold-adaptive traits and effective use of microhabitats (Kirchner 1987). As a result, they constitute a key prey item in avian winter diets across a range of habitats, including coniferous forests (Gunnarsson 1988) agricultural landscapes (Michalko et al. 2022) and urban environments (Chatelain et al. 2025). Interestingly, one species (Rufous-gorgeted Flycatcher) that had a lower trophic position in the winter (Figure 1) also less frequently consumed spiders in the winter compared to the summer (Appendix S1: Figure S1). Therefore, in addition to showing differences in winter diets among individual migrant species, these results also demonstrate the strength of using stable isotope analysis and DNA metabarcoding together, as they complement each other and allow for more effective comparisons of bird diets across seasons and species.

### Dietary flexibility and resource switching in residents

In contrast, high-elevation residents have less seasonal overlap in both isotopic niche and dietary composition which suggests that residents are shifting their diets between functionally distinct taxonomic groups across seasons. As predicted, residents have a lower trophic position in the winter compared with the summer, suggesting a decline in insectivory and increase in frugivory. This was confirmed by DNA metabarcoding, which revealed a 20-35% reduction in key arthropod orders such as such as Lepidoptera, Hemiptera, Diptera and Coleoptera. Additionally, there was a winter increase in the consumption of fruit, particularly from the family Polygonaceae, which fruits (knotweed/*Polygonum* sp.) abundantly at high elevations in our field site in winter. Residents also exhibited greater dietary dispersion in the winter indicating higher individual level variation in diet. This suggests that the lack of preferred arthropod availability leads to individual birds opportunistically eating arthropods available in the winter that they would otherwise avoid. Orders infrequently consumed by birds, such as Symphypleona (springtails), Thysanoptera (thrips) and Neuroptera (lacewings and related species) were in the diets of residents in the winter but not in the summer (Jedlicka et al. 2021, Fernandes et al. 2023).

All resident species had a higher trophic level in the summer compared with the winter, but the degree of the difference varied amongst species (Figure 1). Both species of yuhina (Rufous-vented and Stripe-throated) show the strongest seasonal shift in trophic position and dietary niche. In the winter they showed declines in the consumption of most important arthropod orders (Appendix S1: Figure S1) while an increased consumption of a variety of fruiting plant groups such as, honeysuckle, rubus and knotweeds. Our findings corroborate natural history accounts of the family Zosteropidae (white-eyes and yuhinas), which characterize these species as primarily insectivorous while shifting to berries, seeds, and nectar during periods of insect scarcity (Ali and Ripley 1980, Billerman et al. 2022). Brown-throated Fulvetta, Rufous-capped Babbler and Streak-breasted Scimitar-Babbler showed smaller difference in seasonal trophic position (Figure 1) with the latter also showing substantial overlap in its isotopic niche (Figure 2). All three species form mixed flocks in the winter which may improve their chances of accessing limited arthropod prey (Sridhar et al. 2009). Additionally, the Streak-breasted Scimitar-Babbler with its relatively longer and curved bill may be a more specialized forager allowing it to access arthropods such as beetles and spiders hidden in climatically buffered environments such as beneath tree bark and in leaf litter. Therefore, while they may still consume more plant material in the winter, the relative contribution of arthropods to their diets in either season may not be changing as much compared with the yuhinas. These patterns highlight that, although resident species generally shift their diets across seasons, the extent of this shift varies among species. This variation suggests that behavioural and dietary flexibility both play important roles in determining how different species adapt to and persist in high-elevation environments throughout seasonal changes.

## Conclusion

Among Neotropical frugivores, elevational migrants were more frugivorous than residents and showed a stronger match between dietary preferences and actual diets (Boyle et al. 2011). However, these patterns were documented in only a single season, leaving it unclear whether the observed levels of frugivory persist year-round or are specific to the late breeding season. This distinction is important, as long-distance migrants are known to increase frugivory in autumn as a preparatory strategy for migration (Carter et al. 2024). Ours is the first study to characterise the diets of high elevation breeding birds in both their breeding and non-breeding seasons. Using a combination of stable isotope analysis and faecal DNA metabarcoding our study reveals that elevational migrants are dietary specialists possibly tracking arthropod resources across elevation and season while residents are generalists that can switch their diets from primarily arthropods in summer to include more plant material in winter. However, it can be argued that these patterns may arise independent of food limitation. For instance, if migration is driven by climate tracking (Menon et al. 2023), migrants and residents may simply consume the most readily available resources at that elevation. In this case dietary specialization would be a consequence rather than a cause of migration. However, a recent study across 34 mountain ranges globally found that elevationally migrating species do not seasonally track their thermal niche consistently, highlighting the limited direct effect of temperature on the seasonal distribution of birds in mountains (Somveille et al. 2026). Additionally, while migrants in the Himalayas show patterns consistent with thermal niche tracking they do not fully match their breeding-season temperatures by moving to the lowest available elevations (Menon et al. 2023). This suggests that while climatic constraints influence elevational migration, they are not the sole driver. Dietary resource tracking may also play a significant role where the degree of dietary specialization or flexibility is linked to the variation seen in migratory behaviour. To truly confirm the relative roles of temperature and food resource tracking in shaping migration, food supplementation experiments would be necessary. Although challenging to implement in the short term, supplementing the winter diets of migrants would allow us to assess whether their tendency to migrate is altered, a phenomenon observed in temperate zones where anthropogenic food subsidies have led to shortened or arrested migrations (Van Doren et al. 2021, Marcelino et al. 2023). It is well known that many species of birds tend to be more carnivorous during the breeding season because of higher protein requirements while raising young (Martin 1987). Therefore, in addition to food supplementation experiments, food choice experiments – such as those described by Boyle et al. (2011) – when conducted in both seasons can help confirm if species are actually shifting diets because of availability or because of changing nutritional requirements.

Understanding how temperature and resource availability interact to shape migration has implications for how different species will adapt to environmental change. Tropical montane species are particularly vulnerable to climate-associated warming and shifting phenology. While migrants can potentially move upslope to track thermal requirements, being dietary specialists might make them vulnerable to phenological mismatches, where their arrival no longer aligns with peak arthropod availability. In contrast, the dietary flexibility of residents may provide greater resilience to such uncertainty. Ultimately, this distinction offers crucial insight into which species may be most at risk and how to prioritize conservation efforts in vulnerable mountain biomes.

## Acknowledgements

We are grateful to the Arunachal Pradesh Forest Department – Shergaon Forest Division and the Singchung Bugun Village Community Reserve for permission to conduct fieldwork, and for their continued support of this work. This research was supported by the Kessel Fellowship of the American Ornithological Society and grants from the Science and Engineering Research Board, Ministry of Environment, Forest and Climate Change, Ministry of Education of the Government of India. We would like to thank our field team, Shambu Rai, Mangal Rai, D.K. Pradhan, Binod Munda, Bharat Tamang, Aman Biswakarma, Aiti Thapa and Dema Tamang for help with data collection. We thank Sindhoora P and Vibhas Shevde for help with the stable isotope analysis. We thank Nidhi Yadav, Lakshminarayanan C P, and Awadhesh Pandit for help with the DNA metabarcoding protocols and sequencing. We would also like to thank Kartik Shanker, Robin Vijayan and Sahas Barve for discussions and suggestions at various stages of this manuscript.

