## Appendix S1 for "Resource abundance and dietary specialization predict elevational migration in a hyperdiverse montane bird community"

**Field Supplementary Methods**

Nets were kept open for about six hours during the day for a total of 10-12 days in either season. Nets were checked every 20 minutes and every bird captured was measured and banded with a uniquely numbered aluminium band. We collected blood and faecal samples from all high elevation residents and elevational migrants. Birds were placed in disposable paper bags and kept for up to ten minutes, as soon as they provided a faecal sample they were removed, and blood was extracted via brachial venipuncture and a heparinised capillary tube. Faecal samples were collected from the paper bag (disposed after use) using a sterile polyester swab and placed in Longmire’s solution (Longmire et al. 1997) and stored at room temperature (~ 6 weeks) until taken back to the lab where it was stored at -20°C. Blood samples were stored at room temperature in 70% ethanol.

**DNA metabarcoding supplementary methods**

As part of library preparation for sequencing on the Illumina NovaSeq 6000 Platform, primers were modified with Illumina overhang adapter sequences complementary to the Nextera XT indexing primers. Forward overhang: 5′ TCGTCGGCAGCGTCAGATGTGTAT AAGAGACAG-primer sequence; Reverse overhang: 5′ GTCTCGTGGGCTCGGAGATGTGTATAAGAGACAG‐primer sequence

For libraries prepared for sequencing on the Illumina MiSeq Platform in addition to the Illumina overhang, primers were also modified to include heterogeneity spacers (Naik et al. 2023)

*PCR for Arthropod DNA*

Primer described by Zeale et al. (2011)

Forward (ZBJ-ArtF1c): AGATATTGGAACWTTATATTTTATTTTTGG

Reverse (ZBJ-ArtR2c): WACTAATCAATTWCCAAATCCTCC

PCR composition: 12 μl of QIAGEN Multiplex PCR Master Mix, 1.5 μl of each 2 µM primer, 3 μl of ultra-pure water, and 7 μl of DNA extract

Thermocycler conditions: initial denaturation step at 95 °C for 15 min, followed by 35 cycles of denaturation at 95 °C for 30 s, annealing at 45 °C for 30 s, and extension at 72 °C for 30 s, followed by a final extension step at 72 °C for 10 min

*PCR for Plant DNA*

Primer described by Taberlet et al. (2007)

Forward (trnL g): GGGCAATCCTGAGCCAA

Reverse (trnL h): CCATTGAGTCTCTGCACCTATC

PCR composition: 12 μl of QIAGEN Multiplex PCR Master Mix, 1.5 μl of each 2 µM primer, 4 μl of ultra-pure water, and 6 μl of DNA extract

Thermocycler conditions: initial denaturation step at 95 °C for 10 min, followed by 35 cycles of denaturation at 95 °C for 30 s, annealing at 50 °C for 30 s, and extension at 72 °C for 30 s, followed by a final extension step at 72 °C for 2 min

*Purification*

Samples from both PCR reactions were purified with Agencourt AMPure XP Beads (Beckman Coulter, Fullerton, CA, USA)

For arthropod DNA we used 15 μl (1 ×) of resuspended beads and 15 μl of arthropod DNA

For plant DNA we used 30 μl (2 ×) of resuspended beads and 15 μl of plant DNA

*Indexing PCR*

Purified arthropod and plant PCR products were pooled in equal volume and each sample was indexed with Nextera XT Indexing Kit in a limited cycle PCR

PCR composition: 25 μl of QIAGEN Multiplex PCR Master Mix, 5 μl of each indexing primer and 4 μl of pooled DNA, 11 μl of ultra-pure water

Thermocycler conditions: initial denaturation step at 95 °C for 3 min, followed by 8 cycles of denaturation at 95 °C for 30 s, annealing at 55 °C for 30 s, and extension at 72 °C for 30 s, followed by a final extension step at 72 °C for 5 min.

*Final pooling and purification*

DNA concentrations were quantified using a Qubit Fluorometer (Invitrogen, Darmstadt, Germany), samples were all pooled in equal volume followed by a second purification step using 30 μl (1.5 ×) of resuspended beads and 20 μl of DNA.

The final DNA concentration was quantified and library fragment size (Arthropod: 230-270 bp, Plant: 80-170 bp) was checked on a 2100 Bioanalyzer and Agilent High Sensitivity DNA chip (Agilent Technologies, Santa Clara, USA)

*Sequencing*

Using the pooled libraries, 150-bp paired-end V2 Illumina MiSeq (95 samples; Dec 2023) and S4-2x150bp NovaSeq 6000 (188 samples; Jan 2025) sequencing runs were carried out at the NGS facility at the National Centre for Biological Sciences Bangalore.

**Stable Isotope Supplementary Methods**

*Instruments used*

The blood samples were analysed for δ15 N and δ13C at the Stable Isotope Facility at the Indian Institute of Science Education and Research Pune using an isotope ratio mass spectrometer (Isoprime 100; Isoprime, Elementar) attached to an Vario Pyro cube elemental analyser (Elementar). The plant samples were analysed at the Stable Isotope Facility at the Centre for Earth Sciences in the Indian Institute of Science Bangalore which used a Delta V Advantage IRMS (Thermo Scientific) attached to a Flash 2000 Elemental Analyser.

*Baseline corrected δ^13^ C (Olsson et al. 2009)*

δ^13^ Ccorr = (δ^13^ C_bird_ – δ^13^ C_mean baseline_) / (δ^13^ C_max baseline_ – δ^13^ C_min baseline_)

δ^13^ Ccorr represents the corrected carbon isotope value for an individual bird. δ^13^ C_bird_ represents the measured δ^13^ C value of the individual bird, δ^13^ C_mean baseline_ is the mean δ^13^ C of all sampled plants at an elevation category. δ^13^ C_max baseline_ and δ^13^ C_min baseline_ are the minimum and maximum δ^13^ C of all sampled plants at an elevation category.

**Supplementary Tables and Figures**

Table S1: Table of mean and standard error of baseline (plant) δ^15^N and δ^13^C as well as the minimum and maximum δ^13^C at three elevation categories. These values were used to calculate TP and δ^13^Ccorr

| Elevation | Mean δ^15^N | Mean δ^13^C | Min δ^13^C | Max δ^13^C |
| --- | --- | --- | --- | --- |
| low (600-1500m) | -1.05 (0.31) | -30.30 (0.34) | -27.5 | -34.2 |
| mid (1600-2200m) | -2.58 (0.5) | -30.27 (0.32) | -26.6 | -32.6 |
| high (2300-2800m) | -2.91 (0.37) | -29.63 (0.41) | -25.7 | -32.7 |

Table S2: List of species from which blood and faecal samples were collected. The number of samples processed are represented as n(summer)/n(winter).

|  | Species | Scientific Name | Family | Faecal Samples | Blood Samples |
| --- | --- | --- | --- | --- | --- |
| Resident | Black-faced Laughingthrush | *Trochalopteron affine* | Leiothrichidae | 0/1 | 1/3 |
|  | Brown-throated Fulvetta | *Fulvetta ludlowi* | Paradoxornithidae | 10/14 | 9/15 |
|  | Rufous-capped Babbler | *Cyanoderma ruficeps* | Timaliidae | 6/11 | 9/10 |
|  | Rufous-vented Yuhina | *Yuhina occipitalis* | Zosteropidae | 12/11 | 10/9 |
|  | Streak-breasted Scimitar-Babbler | *Pomatorhinus ruficollis* | Timaliidae | 11/12 | 7/9 |
|  | Stripe-throated Yuhina | *Yuhina gularis* | Zosteropidae | 9/12 | 10/10 |
|  | Brown Parrotbill | *Paradoxornis unicolor* | Paradoxornithidae | 2/2 | 0 |
|  | Black-browed Tit | *Aegithalos iouschistos* | Aegithalidae | 1/2 | 5/4 |
|  | Striated Laughingthrush | *Grammatoptila striata* | Leiothrichidae | 0/1 | 0 |
|  | Yellow-cheeked Tit | *Machlolophus spilonotus* | Paridae | 2/2 | 0 |
| Migrant | Snowy-browed Flycatcher | *Ficedula hyperythra* | Muscicapidae | 3/3 | 0 |
|  | Buff-barred Warbler | *Phylloscopus pulcher* | Phylloscopidae | 12/10 | 10/10 |
|  | Chestnut-headed Tesia | *Cettia castaneocoronata* | Cettiidae | 12/10 | 11/6 |
|  | Rufous-bellied Niltava | *Niltava sundara* | Muscicapidae | 8/8 | 12/11 |
|  | Rufous-gorgeted Flycatcher | *Ficedula strophiata* | Muscicapidae | 12/11 | 8/12 |
|  | Rufous-winged Fulvetta | *Schoeniparus castaneceps* | Pellorneidae | 11/14 | 10/10 |
|  | Whistler's Warbler | *Phylloscopus whistleri* | Phylloscopidae | 14/11 | 13/10 |
|  | White-tailed Robin | *Myiomela leucura* | Muscicapidae | 2/1 | 0 |

Table S3: Parameter estimates of Bayesian linear mixed effects models testing the effect of season and migratory strategy on trophic position of all species pooled (random intercept of species phylogenetic relationships) and individual species.

| Species | MigStrat | Parameter | Estimate | 95% CI | Rhat | Effective Sample Size |
| --- | --- | --- | --- | --- | --- | --- |
| All Species |  | Intercept | 3.61 | [3.18, 4.01] | 1 | 2894 |
|  |  | MigStrat_resident | 0.03 | [-0.36, 0.42] | 1 | 3739 |
|  |  | Season_winter | 0.15 | [-0.02, 0.32] | 1 | 4240 |
|  |  | MigStrat*Season | -0.49 | [-0.73, -0.25] | 1 | 4192 |
|  |  | Phylogenetic SD | 0.38 | [0.21, 0.67] | 1 | 2425 |
| Whistler's Warbler | Migrant | Summer TP (Intercept) | 3.43 | [3.15, 3.73] | 1.001 | 4944 |
|  |  | Winter Shift (Beta) | 0.42 | [-0.01, 0.82] | 1.001 | 5522 |
| Rufous-gorgeted Flycatcher |  | Summer TP (Intercept) | 3.81 | [3.43, 4.16] | 1.001 | 3874 |
|  |  | Winter Shift (Beta) | -0.07 | [-0.51, 0.38] | 1 | 4803 |
| Chestnut-headed Tesia |  | Summer TP (Intercept) | 3.77 | [3.48, 4.08] | 1 | 5116 |
|  |  | Winter Shift (Beta) | 0.39 | [-0.08, 0.84] | 1 | 5251 |
| Rufous-bellied Niltava |  | Summer TP (Intercept) | 4 | [3.72, 4.27] | 1 | 5320 |
|  |  | Winter Shift (Beta) | 0.23 | [-0.15, 0.63] | 1.001 | 5100 |
| Buff-barred Warbler |  | Summer TP (Intercept) | 3.56 | [3.22, 3.89] | 1 | 4861 |
|  |  | Winter Shift (Beta) | -0.03 | [-0.48, 0.41] | 1.001 | 4997 |
| Rufous-winged Fulvetta |  | Summer TP (Intercept) | 3.72 | [3.5, 3.94] | 1 | 4862 |
|  |  | Winter Shift (Beta) | 0.1 | [-0.2, 0.4] | 1 | 5231 |
| Brown-throated Fulvetta | Resident | Summer TP (Intercept) | 3.69 | [3.39, 3.99] | 1.001 | 4855 |
|  |  | Winter Shift (Beta) | -0.09 | [-0.46, 0.29] | 1.001 | 5097 |
| Rufous-capped Babbler |  | Summer TP (Intercept) | 3.89 | [3.58, 4.18] | 1 | 5147 |
|  |  | Winter Shift (Beta) | -0.21 | [-0.61, 0.2] | 1 | 5529 |
| Stripe-throated Yuhina |  | Summer TP (Intercept) | 3.68 | [3.42, 3.92] | 1 | 4559 |
|  |  | Winter Shift (Beta) | -0.74 | [-1.07, -0.38] | 1.001 | 4780 |
| Streak-breasted Scimitar-Babbler |  | Summer TP (Intercept) | 4.01 | [3.68, 4.33] | 1.001 | 4754 |
|  |  | Winter Shift (Beta) | -0.11 | [-0.51, 0.3] | 1.002 | 5506 |
| Rufous-vented Yuhina |  | Summer TP (Intercept) | 3.98 | [3.69, 4.26] | 1.002 | 4748 |
|  |  | Winter Shift (Beta) | -0.62 | [-0.98, -0.23] | 1 | 5032 |

Table S4: Table representing the median Bayesian probability of finding an individual in one season within the same species’ isotopic niche in the other season at an alpha of 0.95.

|  | Species | p(Winter overlaps summer) | p(Summer overlaps Winter) |
| --- | --- | --- | --- |
| Resident | Brown-throated Fulvetta | 14.2 | 30.15 |
|  | Rufous-capped Babbler | 33.2 | 22.6 |
|  | Rufous-vented Yuhina | 13.45 | 27.3 |
|  | Streak-breasted Scimitar-Babbler | 88.45 | 77.6 |
|  | Stripe-throated Yuhina | 13.5 | 65.85 |
| Migrant | Buff-barred Warbler | 87.8 | 52.6 |
|  | Chestnut-headed Tesia | 28.6 | 91.55 |
|  | Rufous-bellied Niltava | 75.9 | 73.55 |
|  | Rufous-gorgeted Flycatcher | 77.3 | 89.15 |
|  | Rufous-winged Fulvetta | 74.2 | 83.55 |
|  | Whistler's Warbler | 57.45 | 36.85 |

Table S5: Parameter estimates of Bayesian logistic mixed effect models testing the effect of season and migratory strategy on frequency of occurrence (FOO) of five arthropod orders and one plant family in the diets of all species pooled (random intercept of species phylogenetic relationships)

| Order/Family | Parameter | Estimate | 95% CI | Rhat | Effective Sample Size |
| --- | --- | --- | --- | --- | --- |
| Araneae | Intercept | -0.31 | [-1.01, 0.43] | 1.001 | 5015 |
|  | Season_winter | 0.82 | [0.13, 1.52] | 1 | 7540 |
|  | MigStrat_resident | 0.39 | [-0.57, 1.31] | 1 | 5656 |
|  | MigStrat*Season | -1.02 | [-2.02, 0.02] | 1 | 6882 |
|  | Phylogenetic SD | 0.44 | [0.03, 1.08] | 1 | 2508 |
| Hemiptera | Intercept | 1.05 | [-0.6, 2.66] | 1.001 | 3076 |
|  | Season_winter | 0.5 | [-0.36, 1.37] | 1 | 6183 |
|  | MigStrat_resident | -0.41 | [-2.32, 1.41] | 1 | 3732 |
|  | MigStrat*Season | -1.45 | [-2.65, -0.26] | 1.001 | 5817 |
|  | Phylogenetic SD | 1.53 | [0.82, 2.61] | 1.001 | 2888 |
| Coleoptera | Intercept | 0.66 | [-0.09, 1.49] | 1.002 | 3744 |
|  | Season_winter | -0.09 | [-0.8, 0.64] | 1.001 | 6065 |
|  | MigStrat_resident | -0.44 | [-1.51, 0.49] | 1.001 | 4141 |
|  | MigStrat*Season | -0.75 | [-1.8, 0.27] | 1.001 | 5421 |
|  | Phylogenetic SD | 0.52 | [0.04, 1.17] | 1.002 | 2147 |
| Lepidoptera | Intercept | 3.46 | [1.65, 5.69] | 1.001 | 3725 |
|  | Season_winter | 0.6 | [-1.16, 2.57] | 1.001 | 5160 |
|  | MigStrat_resident | 0.61 | [-2, 3.31] | 1.002 | 3399 |
|  | MigStrat*Season | -3.81 | [-6.5, -1.48] | 1.001 | 4136 |
|  | Phylogenetic SD | 1.4 | [0.3, 3.11] | 1.001 | 2292 |
| Diptera | Intercept | 4.03 | [1.87, 6.57] | 1 | 3581 |
|  | Season_winter | -1.95 | [-3.89, -0.4] | 1.001 | 4122 |
|  | MigStrat_resident | -2.01 | [-4.69, 0.5] | 1 | 3772 |
|  | MigStrat*Season | 0.84 | [-1.02, 3.03] | 1.001 | 4129 |
|  | Phylogenetic SD | 1.67 | [0.72, 3.17] | 1.001 | 2950 |
| Polygonaceae | Intercept | -0.97 | [-2.35, 0.38] | 1 | 3686 |
|  | Season_winter | -0.92 | [-1.8, -0.09] | 1 | 7098 |
|  | MigStrat_resident | -1.46 | [-3.44, 0.1] | 1 | 3777 |
|  | MigStrat*Season | 2.97 | [1.66, 4.38] | 1 | 5838 |
|  | Phylogenetic SD | 1.13 | [0.17, 2.60] | 1 | 1371 |

Table S6: Table summarising the frequency of occurrence (FOO) of a few plant families in the diets of residents and migrants in either season. We have included families with the four highest FOOs in the diets of residents and migrants in each season.

|  | Family | Summer | Winter |
| --- | --- | --- | --- |
| Resident | Acanthaceae | 0.02 | 0.01 |
|  | Caprifoliaceae | 0.06 | 0.16 |
|  | Ericaceae | 0.68 | 0.15 |
|  | Fagaceae | 0.11 | 0.04 |
|  | Malvaceae | 0.00 | 0.01 |
|  | Pinaceae | 0.09 | 0.01 |
|  | Poaceae | 0.15 | 0.04 |
|  | Polygonaceae | 0.11 | 0.44 |
|  | Rosaceae | 0.19 | 0.22 |
| Migrant | Acanthaceae | 0.00 | 0.18 |
|  | Caprifoliaceae | 0.08 | 0.00 |
|  | Ericaceae | 0.57 | 0.02 |
|  | Fagaceae | 0.14 | 0.09 |
|  | Malvaceae | 0.00 | 0.15 |
|  | Pinaceae | 0.22 | 0.00 |
|  | Poaceae | 0.08 | 0.05 |
|  | Polygonaceae | 0.30 | 0.15 |
|  | Rosaceae | 0.30 | 0.21 |

Table S7: Results of PERMANOVA test with 999 permutations comparing seasonal differences in diet composition in terms of the arthropod and plant orders. Degrees of freedom (df) reported as “numerator df/denominator df”.

|  | Species | Pseudo F | df | p value |
| --- | --- | --- | --- | --- |
| Residents | Brown-throated Fulvetta | 1.41 | 1/22 | 0.12 |
|  | Rufous-capped Babbler | 1.96 | 1/15 | 0.03 |
|  | Rufous-vented Yuhina | 9.24 | 1/21 | 0.001 |
|  | Streak-breasted Scimitar-Babbler | 3.33 | 1/21 | 0.001 |
|  | Stripe-throated Yuhina | 3.51 | 1/19 | 0.001 |
| Migrant | Buff-barred Warbler | 3.29 | 1/18 | 0.001 |
|  | Chestnut-headed Tesia | 1.7 | 1/21 | 0.01 |
|  | Rufous-bellied Niltava | 2.33 | 1/15 | 0.001 |
|  | Rufous-gorgeted Flycatcher | 1.18 | 1/21 | 0.21 |
|  | Rufous-winged Fulvetta | 2.51 | 1/23 | 0.001 |
|  | Whistler's Warbler | 2.02 | 1/23 | 0.005 |

Table S8: Results of PERMDISP test with 999 permutations comparing seasonal differences in dietary dispersion in terms of the arthropod and plant orders. Degrees of freedom (df) reported as “numerator df/denominator df”.

|  | Species | Pseudo F | df | p value |
| --- | --- | --- | --- | --- |
| Residents | Brown-throated Fulvetta | 4.53 | 1/22 | 0.04 |
|  | Rufous-capped Babbler | 0.86 | 1/15 | 0.38 |
|  | Rufous-vented Yuhina | 178.3 | 1/21 | <0.001 |
|  | Streak-breasted Scimitar-Babbler | 0 | 1/21 | 0.99 |
|  | Stripe-throated Yuhina | 9.51 | 1/19 | 0.01 |
| Migrant | Buff-barred Warbler | 1.27 | 1/18 | 0.31 |
|  | Chestnut-headed Tesia | 0.89 | 1/21 | 0.35 |
|  | Rufous-bellied Niltava | 0.4 | 1/15 | 0.54 |
|  | Rufous-gorgeted Flycatcher | 2.23 | 1/21 | 0.15 |
|  | Rufous-winged Fulvetta | 0.96 | 1/23 | 0.34 |
|  | Whistler's Warbler | 0.14 | 1/23 | 0.72 |

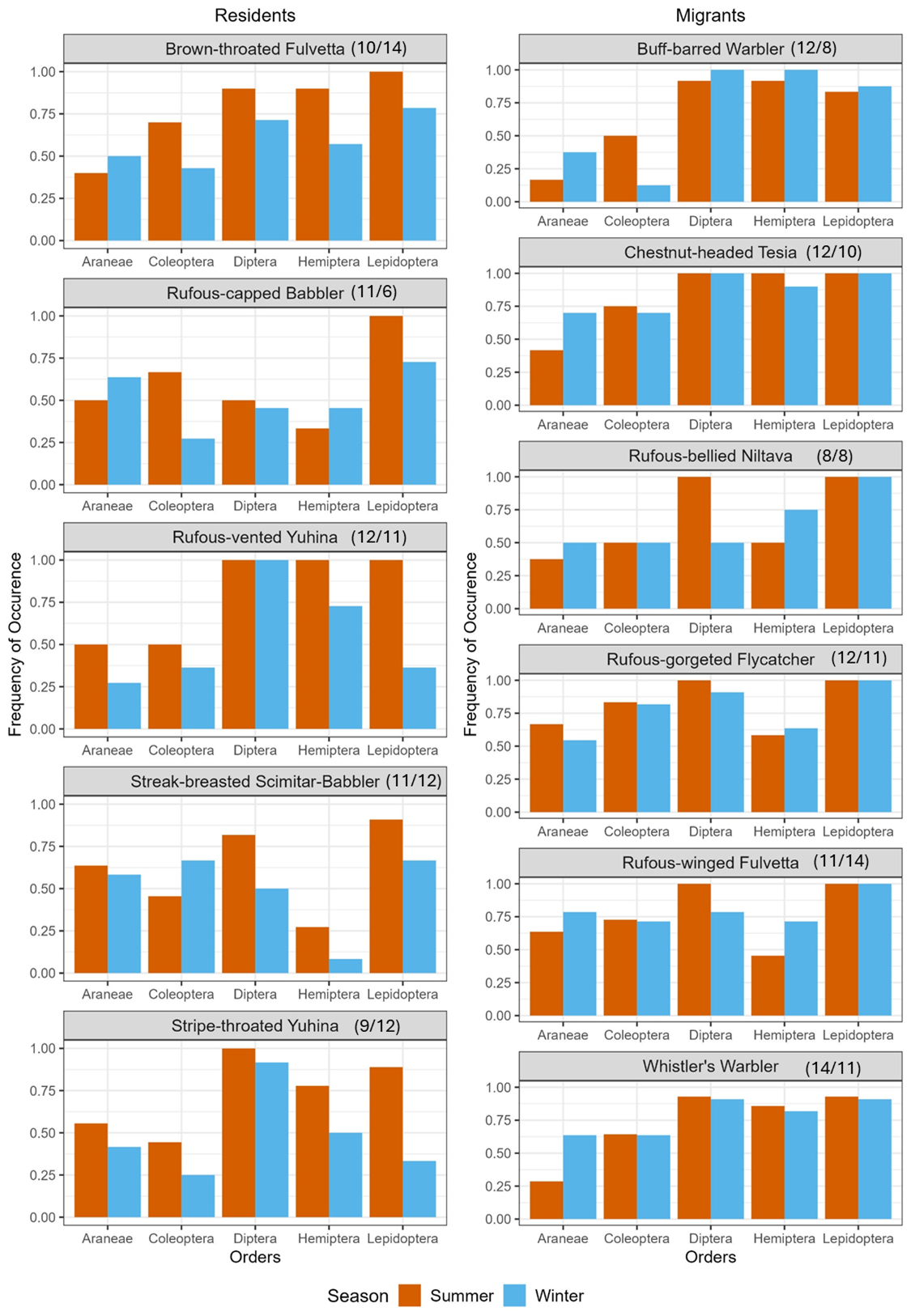

Figure S1: Bar plots comparing the frequency of occurrence of the five most consumed arthropod orders in the diets of individual species of high-elevation residents and migrants across seasons. Asterisks represent a significant two-sample proportion test. Sample sizes in the summer/winter given in parenthesis at the top of each plot.

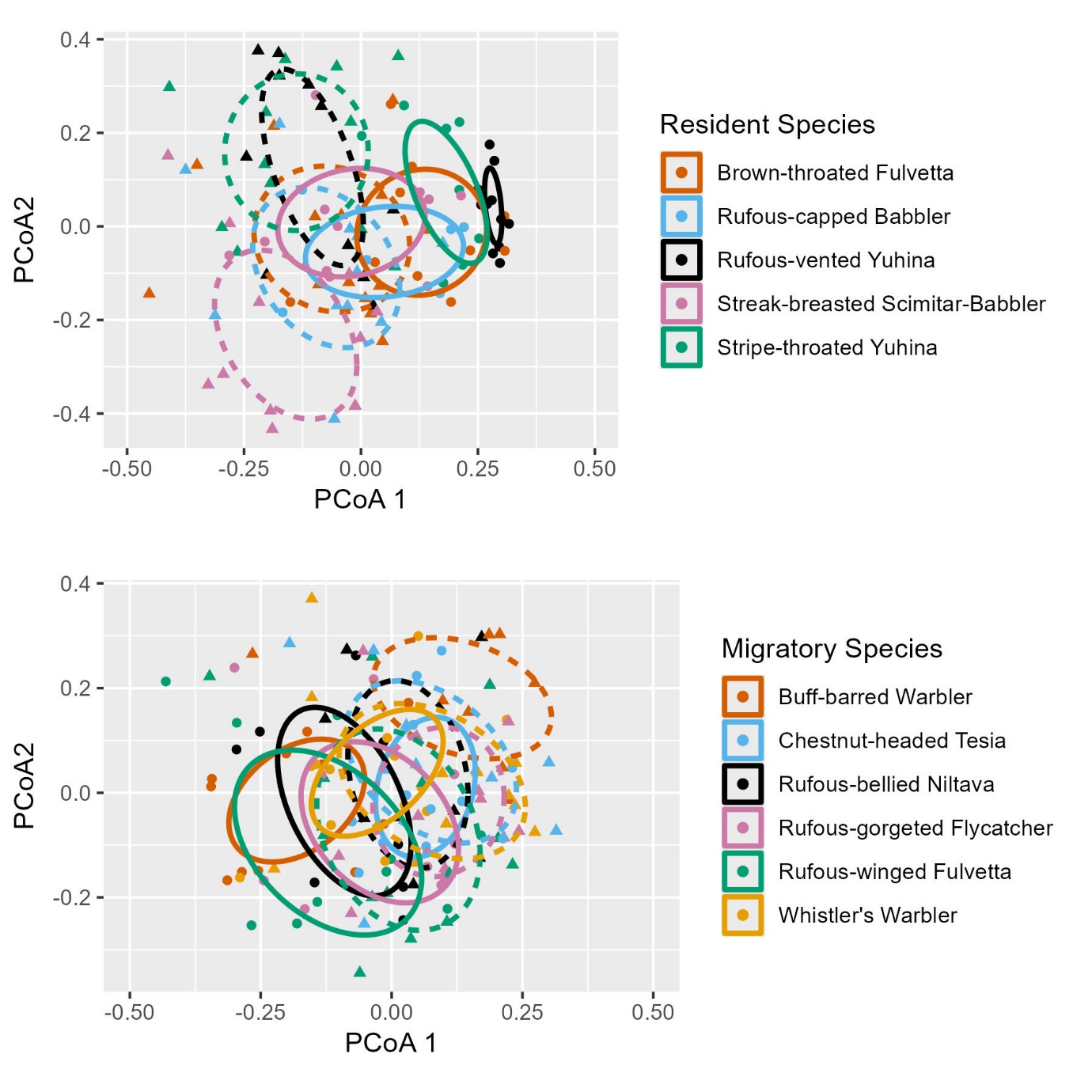

Figure S2: Principal Coordinate Analysis of differences in dietary composition of individual migratory and high elevation resident species across seasons. Circles represent summer diet and triangles represent winter diet. Ellipses denote the standard error of the centroid diet for each species in a particular season at 95% confidence with solid lines indicating summer diets and dotted lines indicating winter diets
